# Herpesvirus infection reduces Pol II occupancy of host promoters but spares viral promoters

**DOI:** 10.1101/585984

**Authors:** Ella N Hartenian, Britt A Glaunsinger

## Abstract

In mammalian cells, widespread acceleration of cytoplasmic mRNA degradation is linked to impaired RNA polymerase II (Pol II) transcription. This mRNA decay-induced transcriptional repression occurs during infection with gammaherpesviruses including Kaposi’s sarcoma-associated herpesvirus (KSHV) and murine gammaherpesvirus 68 (MHV68), which encode an mRNA endonuclease that initiates widespread RNA decay. Here, we show that MHV68-induced mRNA decay leads to a genome-wide reduction of Pol II occupancy at mammalian promoters. Viral genes, despite the fact that they require Pol II for transcription, escape this transcriptional repression. Protection is not governed by viral promoter sequences; instead, location on the viral genome is both necessary and sufficient to escape the transcriptional repression effects of mRNA decay. We hypothesize that the ability to escape from transcriptional repression is linked to the localization of viral DNA in replication compartments, providing a means for these viruses to counteract decay-induced viral transcript loss.

## Introduction

Regulating messenger RNA (mRNA) abundance is of central importance during both cellular homeostasis and disease. While it is intuitive that transcriptional changes in the nucleus impact RNA levels in the cytoplasm, evidence in both yeast and mammalian cells now indicates that the converse is also true: altered cytoplasmic mRNA decay rates can broadly impact transcription by RNA polymerase II (Pol II). In yeast, stabilizing the cytoplasmic mRNA pool by removing mRNA decay factors such as Xrn1 causes a compensatory decrease in Pol II transcription (1-5). Furthermore, reducing or slowing Pol II transcription leads to an increase in overall mRNA stability (1), supporting a “buffering” model in which yeast can compensate for widespread perturbations to mRNA abundance by alternatively changing rates of transcription.

Broad changes in mRNA abundance are often triggered in mammalian cells during viral infection. Numerous viruses, including alphaherpesviruses, gammaherpesviruses, SARS coronavirus and influenza A virus drive accelerated mRNA decay during infection by expressing mRNA specific endoribonucleases and/or activating host nucleases. This decay contributes to viral immune evasion, increases the availability of host translation machinery, and facilitates temporal viral gene regulation (6-10). Viral endonucleases target host and viral mRNAs for cleavage, whereupon cellular exonucleases degrade the resulting mRNA fragments (11-15). This strategy accelerates basal RNA decay by circumventing the typically rate limiting steps of deadenylation and decapping (16-20).

Although yeast and mammals share key proteins involved in cytoplasmic mRNA decay and transcription, studies with RNA-decay inducing viruses in mammals have revealed that unlike the yeast pathway, accelerated mRNA decay leads to an extensive shutoff of cellular gene expression (21,22). Infection with the gammaherpesviruses KSHV or MHV68 causes reduced Pol II recruitment at host promoters in a manner dependent on mRNA cleavage by the viral endonucleases SOX or muSOX. Thus, accelerated mRNA decay is not broadly counteracted by increased transcription-based repopulation, but instead leads to a more extensive shutdown of cellular gene expression. However, similar to yeast, the mammalian connection between decay and transcription requires the activity of cellular exonucleases such as Xrn1, which degrade the viral endonuclease cleavage products (16,21). Accelerated Xrn1-mediated degradation causes trafficking of released RNA binding proteins (RBP) from the cytoplasm to the nucleus, which may provide a means for the cell to detect and respond to increased RNA turnover (22).

During MHV68 infection, in addition to host mRNA, cytoplasmic viral mRNA is susceptible to cleavage by muSOX (23). However, measurements of nascent transcript production from a subset of viral promoters suggested that viral genes are robustly transcribed during the stage of infection when host transcription is reduced (24). An outstanding question is how these viral promoters escape mRNA decay-induced transcriptional repression despite being transcribed by RNA Pol II. Here, we measured Pol II occupancy across the mouse and MHV68 genomes in infected cells to more comprehensively define how accelerated mRNA decay impacts polymerase occupancy. Our data demonstrate a genome-wide reduction in Pol II recruitment at host promoters under conditions of increased mRNA decay. In contrast, Pol II occupancy of the viral genome appears broadly resistant to the effects of mRNA degradation. This protection is not conferred by viral promoter sequences; instead, location on the replicating viral genome is both necessary and sufficient to escape the transcriptional effects of mRNA decay. We propose a model in which DNA amplified in viral replication compartments, unlike the cellular chromatin, is immune to transcriptional repression during accelerated cytoplasmic mRNA decay. Thus, while both viral and cellular mRNA pools are susceptible to cytoplasmic degradation, transcriptional repopulation selectively counteracts this loss for viral genes.

## Results

### Accelerated RNA decay broadly reduces Pol II occupancy

We previously reported that Pol II occupancy at several individual mammalian promoters was significantly reduced during accelerated cytoplasmic mRNA decay (22,24). To more comprehensively assess the extent of mRNA decay-induced transcriptional repression, we evaluated global Pol II occupancy by chromatin immunoprecipitation and deep sequencing (ChIP-seq) in mock versus MHV68 infected MC57G mouse fibroblasts at 24 hours post infection (hpi). As a control, we also infected cells with a version of MHV68 containing the point mutation R443I in the viral muSOX endonuclease gene, which reduces its mRNA cleavage activity. R443I retains a similar lytic cycle and infectious capability to wildtype (WT) MHV68 *in vivo* and in vitro its infectious capability is similar to MHV68 in mouse fibroblasts (6,23). MHV68-infected cells displayed promoter-proximal loss of Pol II at 86% of loci compared to mock-infected cells, averaged across two biological replicates and subtracting input signal (Fig 1A and S1 Fig A). Loss of Pol II occupancy was primarily due to muSOX-induced mRNA decay, as this signal loss either did not occur or was reduced at 67% of these loci during infection with R443I MHV68 (Fig 1A and S1 Fig A). Sixty-three genes, or 1.3% of loci, showed higher Pol II occupancy in WT infection as compared to mock infection, although these did not share any common GO terms (S1 Table). A representative genome browser view shows reduced Pol II signal throughout *Srsf2* in the MHV68 condition of two replicate ChIP experiments. We independently validated the ChIP-seq results by ChIP-qPCR at the *Rplp0* and *Fus* promoters, both of which showed a significant reduction of Pol II occupancy in cells infected with WT but not R443I MHV68 (Fig 1C), as did a validation using a second antibody that recognizes the N-terminus of the major subunit of Pol II, Rpb1 (Fig1D). The genome-wide reduction in poised Pol II in WT but not R443I infected cells was also evident when the sequencing reads from positions −2000 to +4000 around the transcription start sites (TSS) were plotted as a histogram (Fig 1G and S1 Fig B). The variability between replicates in R443I recovery (Fig 1B) is likely due to differences in the extent of host shutoff reduction between replicates due to R443I reducing muSOX activity while not being a catalytic mutant.

**Fig 1:**
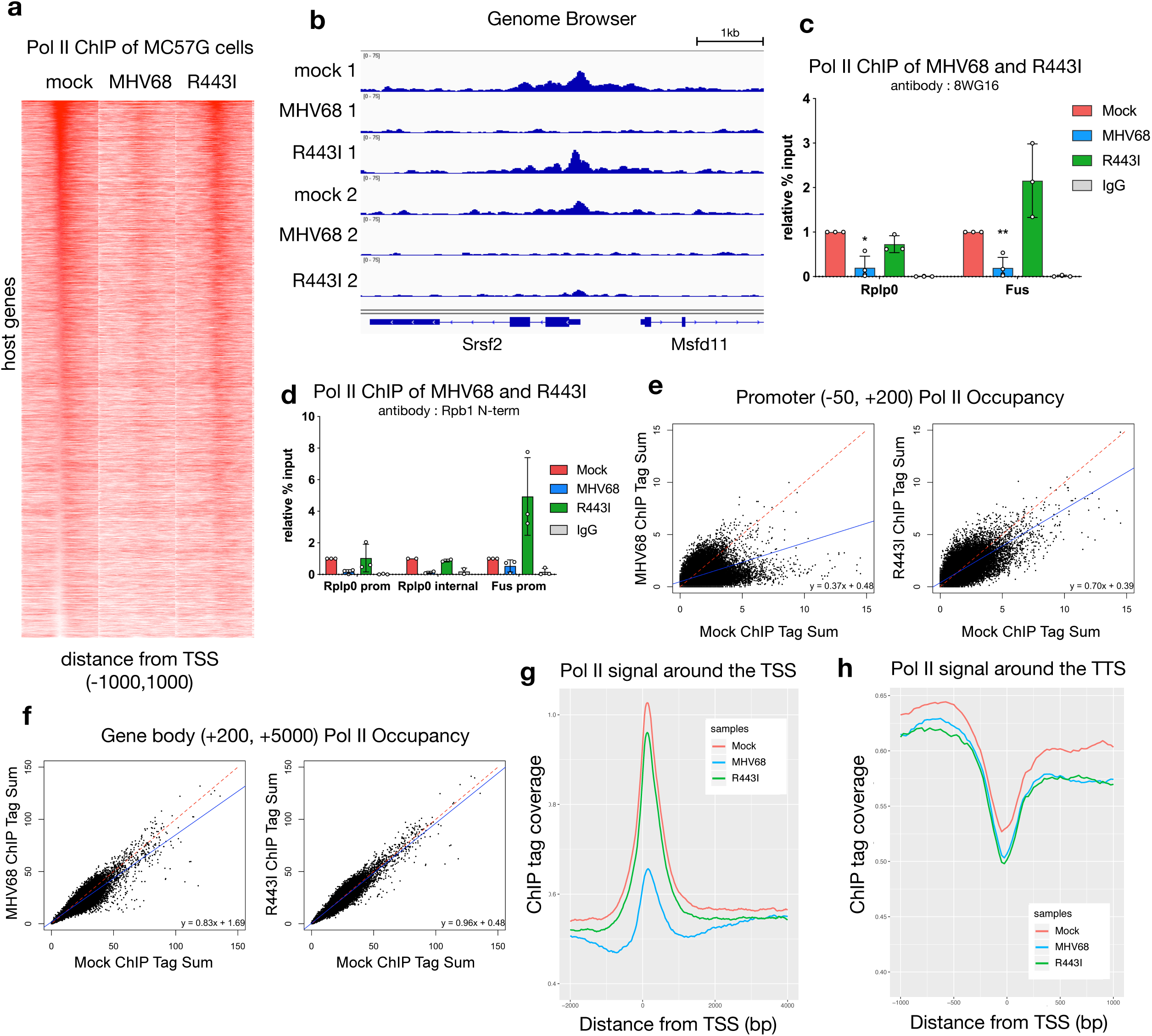
Promoter-proximal Pol II recruitment to mammalian genes is RNA-decay dependent. (A) Pol II ChIP-seq signal profiles of host genes in mock-infected, MHV68-infected and R443I-infected MC57G cells. Each row of the heat map displays Pol II occupancy of one gene from – 1000 to +1000 in 25 bp bins. Genes are ranked by the Pol II-transcription start site (TSS) proximal signal in mock infected cells. (B) ChIP-qPCR validation of Pol II occupancy at the *Rplp0* and *Fus* promoters. Pol II ChIP was performed on mock, MHV68 WT or MHV68 R443I infected MC57G cells and Pol II levels were assayed near the TSS of two repressed host genes during MHV68 infection from the ChIP-seq data. (* p < 0.05, ** p < 0.01, students t-test) (C) Sequence tags were plotted as a histogram with 25 bp bins for −2000 to +4000 around the TSS. Mock (red), MHV68 (blue) and MHV68 R443I (green) traces are shown along with their input controls. (D) Scatterplot of Pol II occupancy of promoters, averaging the sum of ChIP-seq tags from −50 to +200 across two replicate experiments with inputs subtracted out. Mock is compared to WT MHV68 infection on the left and to MHV68 R443I infection on the right. Linear regression lines are plotted in blue and equations are provided. A y=x line is plotted for reference in red. (E) Scatterplot of Pol II occupancy of gene bodies, averaging the sum of ChIP-seq tags from −50 to +200 across two replicate experiments with inputs subtracted out. Data as described for (D) (E) Pol II transcription termination is not dependent on RNA decay. Sequence tags were plotted as a histogram in 25 bp bins for transcription termination sequence (TTS) proximal Pol II for −1000 to +1000 around the TTS with the same color scheme as (C).

To assess the stage of transcription impacted by RNA decay, we compared the amount of Pol II in promoter regions (−50 to +200) and the amount of Pol II in gene bodies (+200 to +5000) between mock and infected samples. Consistent with the above analyses, MHV68 infected cells contained markedly less promoter proximal Pol II than uninfected cells, and this signal was partially recovered in the R443I infection (Fig 1E). Genes with impaired Pol II promoter binding during MHV68 infection also had less Pol II within the gene body compared to mock infected cells, which again was not observed during R443I infection (Fig 1F). Notably, the difference in regression line slopes between MHV68 and mock infected cells was clearly more marked when comparing Pol II promoter occupancy, suggesting that the primary transcriptional defect is at the stage of initiation. This is consistent with data from previous studies that found no defect in serine 2 phosphorylation of elongating Pol II and no consistent change to Pol II elongation during MHV68 infection (22,24).

Finally, to assess if RNA decay affected transcription termination, we plotted sequencing reads between 1kb upstream and 1kb downstream of the transcription termination site (TTS) as a histogram (Fig 1H and S1 Fig D). TTS-proximal Pol II shows no defect in termination in either infection condition, suggesting Pol II removal from host transcripts is not impacted by MHV68 infection or by RNA decay.

### Pol II recruitment to viral genes is not affected by RNA decay

Herpesviral genes are transcribed in the nucleus by mammalian Pol II, and thus could be similarly subjected to the mRNA decay-induced transcriptional repression observed across the host genome. We therefore analyzed Pol II occupancy of the viral genome in WT MHV68 and R443I infected cells to determine how viral genes respond to RNA decay. The average Pol II signature across 82 viral ORFs between −500 and +500 around the TSS was comparable in cells infected with WT or R443I MHV68, with a slightly higher signal in the R443I infection (Fig 2A). Indeed, there was a robust and widespread Pol II signal in both infection conditions across the viral genome (Fig 2B). Thus, herpesviral genes appear to escape mRNA decay-induced transcriptional repression.

**Fig 2:**
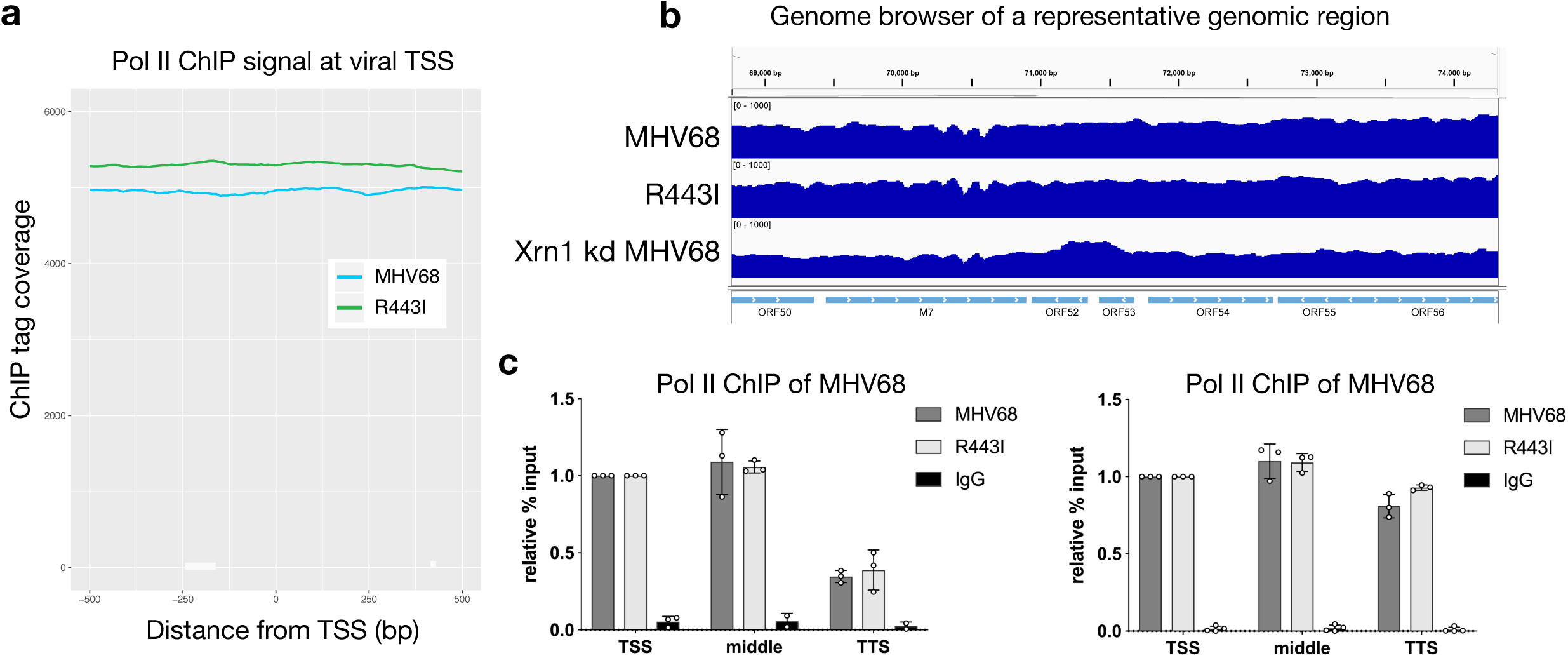
Viral genes are not susceptible to RNA decay dependent Pol II repression. (A) Viral gene Pol II recruitment is independent of RNA decay. Sequence tags were plotted as a histogram in 25 bp bins for −500 to +500 around the TSS. MHV68 WT (blue) and R443I (green) traces are shown from infected MC57G cells. (B) Pol II ChIP-seq coverage across 5.5 kb of the MHV68 genome for MHV68 WT and R443I in wild type (WT) cells and MHV68 infection in Xrn1 kd cells. Alignment files were converted to tdf format and visualized in the IGV. (C) Pol II ChIP qPCR signal is similar across gene bodies. Three regions of the ORF54 and ORF37 gene were assayed by Pol II ChIP-qPCR during MHV68 WT or R443I infection to assess the relative levels of Pol II across the gene.

In contrast to host genes, we did not observe peaks of promoter-proximal Pol II on the viral genome in our ChIP-seq data (Fig 2A and B). Pol II ChIP qPCR across various regions of ORF54 and ORF37 also showed a similar Pol II signal throughout each gene body and at their promoter. We did note a decrease in signal near the TTS of ORF54, however this was true for both WT and R443I infections (Fig 2C). We hypothesize that pervasive transcription of the viral genome at 24 hpi, compounded by the presence of TSSs on both Watson and Crick strands, likely contributes to this phenotype.

### PABPC is not excluded from viral replication compartments

We next considered how the viral genome maintains high levels of Pol II occupancy during accelerated mRNA decay. During replication, viral DNA is transcribed within viral replication compartments (RCs) in the nucleus, which exclude host DNA and thus stain poorly with DAPI (25-27). Although they are not membrane bound, these compartments selectively enrich for factors required for viral replication (such as the viral processivity factor ORF59) and gene expression (such as Pol II) (28-30). Virus-induced mRNA decay causes increased trafficking of cytoplasmic poly(A) binding protein (PABPC) from the cytoplasm to the nucleus, which we have recently linked to transcriptional repression (22,31). To determine whether replication compartments selectively exclude nuclear PABPC as a means of avoiding transcriptional repression, we monitored its localization in cells lytically infected with MHV68 or KSHV. As expected, the majority of infected cells displayed increased nuclear PABPC compared to unreactivated or uninfected cells as measured by immunofluorescence (IF). We further saw that PABPC was not excluded from replication compartments (Fig 3A, B and S2 Fig A). To determine PABPC localization relative to RCs, we first identified cells with RCs by looking for regions of Pol II or ORF59 staining that excluded DAPI. We then quantified nuclear PABPC pixel intensity, and counted cells as having an overlapping RC/PABPC signal when the PABPC pixel intensity was at least 2x that of cells without nuclear PABPC from the same image. Indeed, 85 percent of MHV68 infected cells showed overlapping PABPC/Pol II signal and 90 percent of reactivated iSLKs showed an overlapping PABPC/ORF59 signal (Fig 3C). Thus, PABPC exclusion from RCs is unlikely to underlie viral promoter escape.

**Fig 3:**
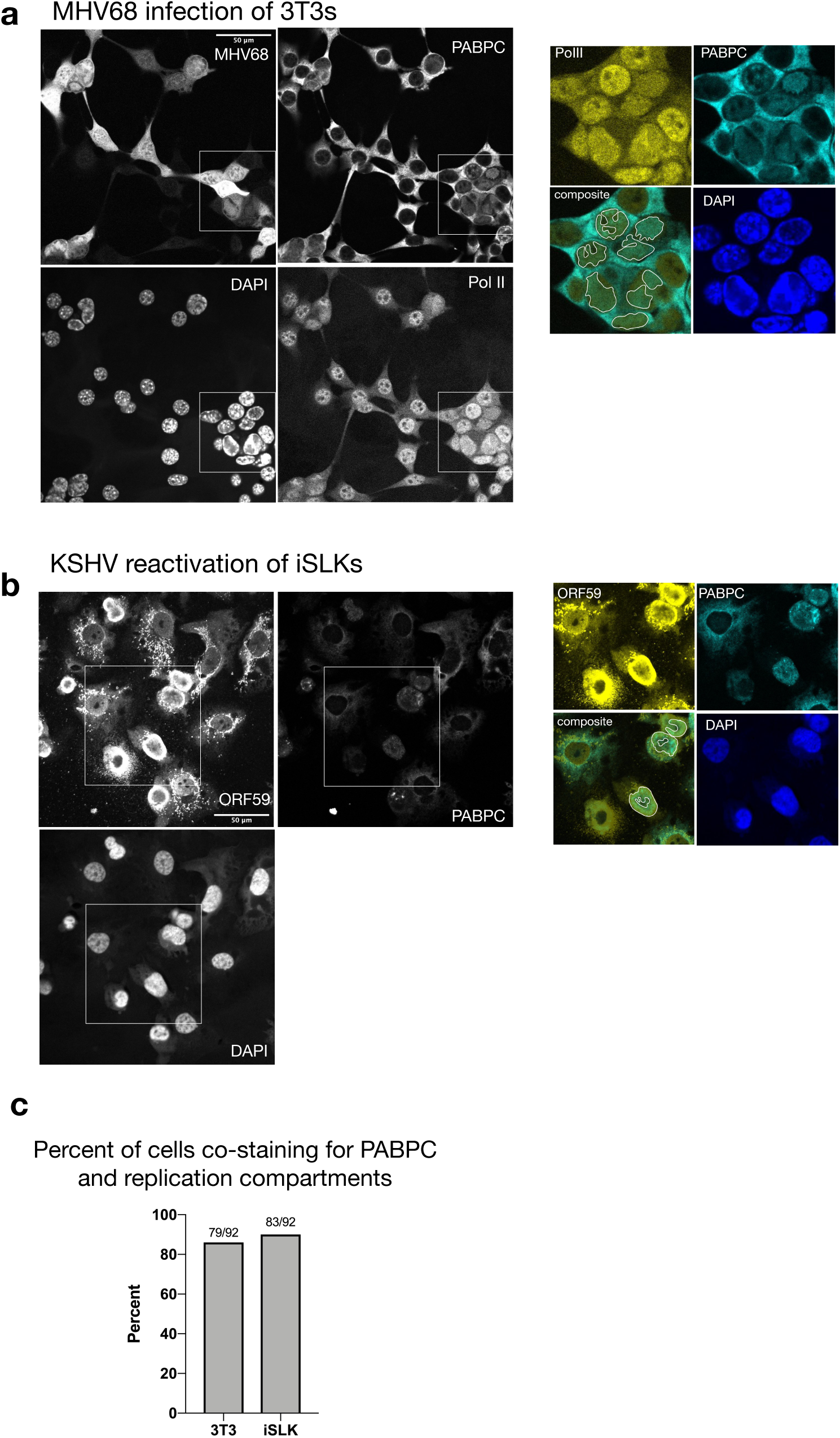
PABPC is not excluded from replication compartments. (A) Imunofluoresence (IF) was performed on MHV68 infected 3T3 cells at 27 hpi and stained with PABPC, Pol II and DAPI. The MHV68 genome contains GFP, which served as a marker of infection. Cells with RCs are denoted with red arrows. RCs were identified in cell nuclei with PolII staining overlapping regions that are DAPI negative. The inset shows a merge of Pol II and PABPC staining for several cells that co-stain for both proteins in RCs. (B) IF was performed on KSHV-positive iSLK cells reactivated for 48 h and stained with antibodies against PABPC, ORF59 and DAPI. Cells with RCs are denoted with arrows and were identified in cell nuclei with ORF59 staining that overlaps with DAPI negative regions. The inset shows a merge of ORF59 and PABPC staining for several cells that co-stain for both proteins in RCs. (C) Percent of RC containing cells that overlap with PABPC signal. Fractions of cells with PABPC signal in RCs over total number of cells counted with RCs are displayed.

### Viral promoters fail to escape transcriptional repression outside of the viral genome

We then tested whether viral promoter sequence elements were sufficient to enable escape by querying whether they conferred resistance to transcriptional repression upon relocation from the viral genome into the host DNA. We inserted the MHV68 late promoter M7 or the KSHV LANA promoter upstream of a puromycin resistance cassette into 293T cell chromatin using either lentiviral transduction or random plasmid DNA integration, respectively. We induced mRNA decay and host transcriptional repression in these cells by expression of MHV68 muSOX or the herpes simplex virus type 1 (HSV-1) endonuclease vhs. Similar to the host promoters *Gapdh* and *Rplp0*, Pol II recruitment to both integrated viral promoters was repressed upon infection with MHV68 but not with R443I (Fig 4A and 5B).

**Fig 4:**
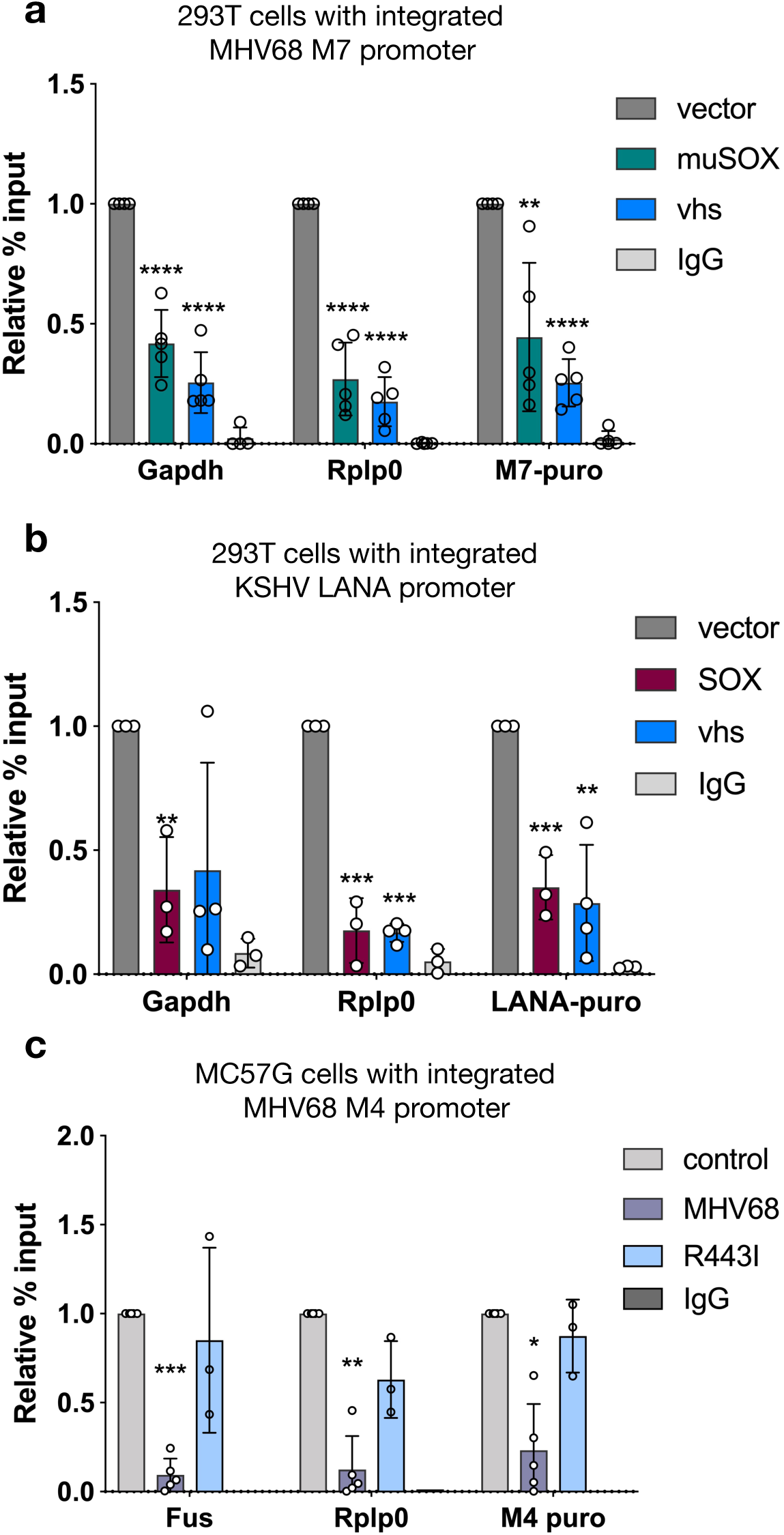
Viral promoter sequences are insufficient to escape transcriptional regulation. (A) The MHV68 M7 viral promoter driving puromycin resistance was lentivirally integrated into 293T cells. 24 h post transfection of muSOX or VHS, Pol II-ChIP qPCR was used to measure Pol II levels at two host promoters (*Gapdh, Rplp0*) and the integrated M7 promoter. (* p < 0.05, ** p < 0.01, *** p <0.001, **** p<0.0001 students t-test) (B) The KSHV LANA promoter driving puromycin resistance was integrated into 293T cells by random incorporation. Pol II-ChIP qPCR was used to measure Pol II levels 24 h post SOX or VHS transfection as described above. (C) The MHV68 M4 viral promoter driving puromycin resistance was lentivirally integrated into MC57G cells. The cells were then infected with WT MHV68 or R443I at a MOI of 5 and Pol II levels at the indicated promoters were assayed by Pol II ChIP-qPCR.

**Fig 5:**
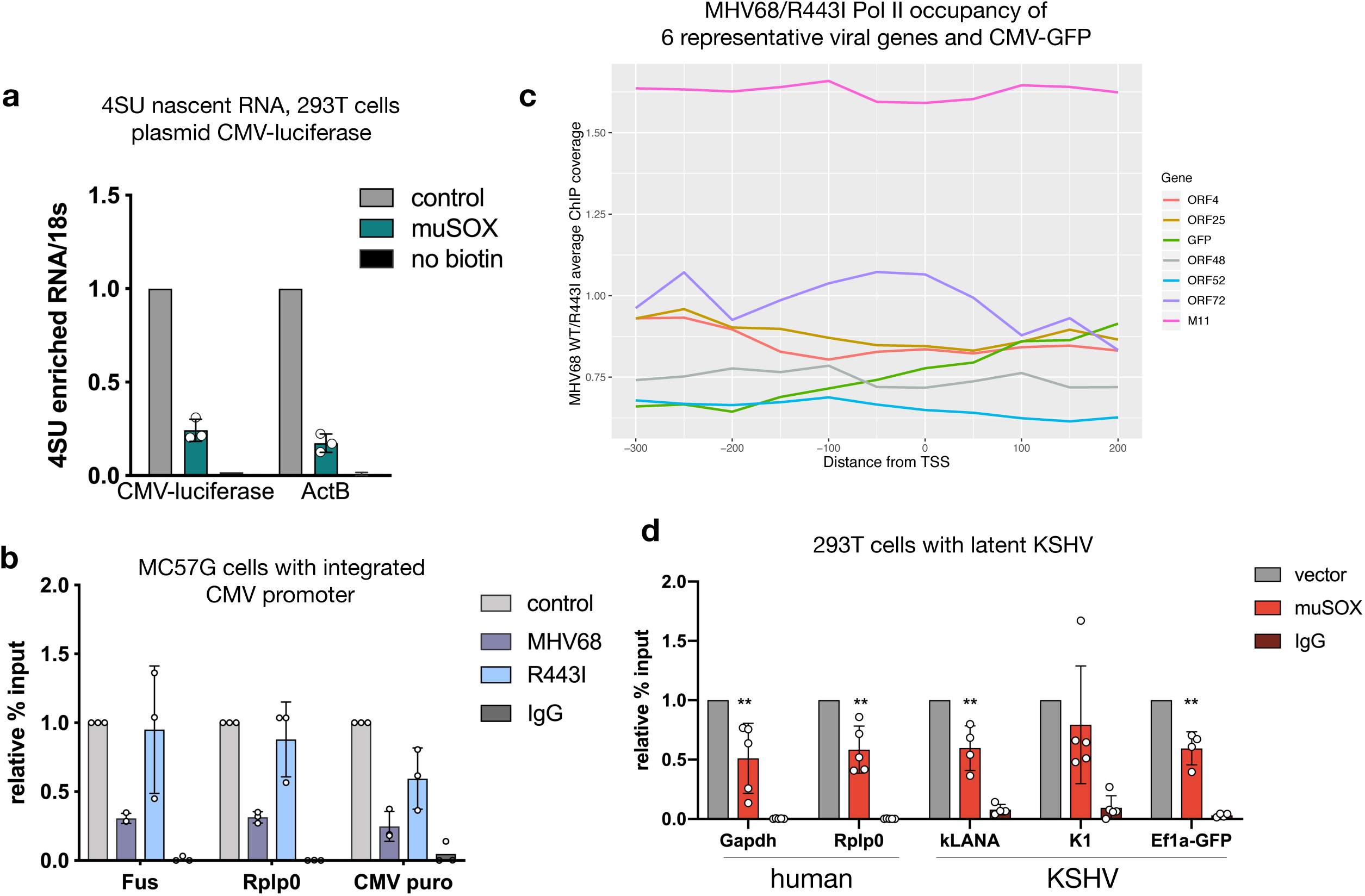
Non-MHV68 promoters escape repression on the viral genome. (A) 4SU incorporated RNA levels of luciferase driven by CMV in muSOX and vector-control expressing 293T cells. Cells were separated using the co-expressed Thy1.1 marker to ensure pure populations of transfected cells(22). (B) The CMV promoter driving puromycin resistance was lentivirally integrated into MC57G cells. The cells were then infected with WT MHV68 or R443I at an MOI of 5 and Pol II levels at the indicated promoters were assayed by Pol II ChIP-qPCR. (C) ChIP-seq traces comparing WT MHV68 to R443I coverage from −300 to +200 around the TSS in 25 bp bins, averaged from two biological replicates of infected MC57G cells. Six representative viral genes and CMV-GFP are shown. (D) 293T cells were infected with KSHV and then transfected with muSOX twice over a 24 h period to induce accelerated decay. Pol II levels at the indicated KSHV and human promoters were measured by ChIP-qPCR. (** p < 0.01, students t-test)

To confirm that viral promoters integrated into the host genome were also repressed in the context of infection, we infected MC57G cells containing an integrated MHV68 M4 promoter-driven puromycin cassette with MHV68. Indeed, the M4 promoter was repressed to a similar degree as host *Fus* and *Rplp0* promoters during infection with WT MHV68 but not the mRNA decay-deficient MHV68 R443I (Fig 4C). In summary, these data argue against the presence of protective elements within viral promoters, and instead suggest that the chromatin context or location on the viral genome underlies escape from mRNA decay-induced transcriptional repression.

### Non-MHV68 promoters escape repression on the viral genome

A prediction from the above results is that non-MHV68 promoters would gain protection from transcriptional repression when present on the replicating viral genome. To test this, we measured the Pol II occupancy and transcriptional potential of the cytomegalovirus (CMV) promoter, which drives green fluorescent protein (GFP) expression on the MHV68 genome. Unlike alpha- and gammaherpesviruses, the betaherpesvirus CMV does not accelerate mRNA decay. Furthermore, in either a plasmid context or when integrated into 293T cells, the CMV promoter is strongly repressed during muSOX-induced mRNA decay. This is seen by 4SU incorporation measuring nascent RNA production – similar to the control *ActB* gene – and by Pol II recruitment, similar to the control *Fus* and *Rplp0* promoters (Fig 5A and B). However, in the context of the replicating MHV68 genome, the CMV-GFP promoter had a Pol II ChIP-seq signal during both MHV68 WT and R443I infection that was not markedly different from several other viral genes (ORFs 4, 25, 48, 52, 72, M11) (Fig 5C).

Finally, we evaluated whether the lytic stage of infection was necessary to confer escape from transcriptional repression, or whether some other feature of the non-replicating viral genome was important. To this end, we examined Pol II occupancy at promoters of several genes expressed from the KSHV genome during latent infection in 293T cells, when the viral DNA is maintained as an episome but not amplified in replication compartments. Latent infection does not promote mRNA turnover, and thus we transfected muSOX into these cells to stimulate mRNA turnover in the absence of lytic replication. Similar to the host genes *Gapdh* and *Rplp0*, we observed an RNA decay-induced repression of Pol II occupancy at the latent viral promoter LANA while the viral K1 promoter was repressed in 4 out of 5 experiments (Fig 5D). We also observed repression of the human *Ef1α* promoter, which is present on the KSHV viral episome as a driver of the GFP reporter gene (Fig 5D). In sum, these observations indicate that the viral genome per se does not confer protection from transcriptional repression; instead, features linked to lytic genome amplification in replication compartments facilitate robust Pol II recruitment under conditions of accelerated mRNA turnover.

## Discussion

Here, we demonstrate that virus-induced mRNA decay broadly decreases Pol II recruitment to mammalian promoters, extending prior observations made with individual cellular genes to the genomic scale. In contrast, promoters on the replicating viral genome recruit Pol II with similar efficiency during normal or accelerated mRNA decay, indicating that they escape transcriptional inhibition. This protection is lost upon relocation of the viral promoters into the host genome or when the viral genome is in a latent state, emphasizing the importance of DNA context and replication state, rather than promoter sequence, in determining the transcriptional response to accelerated mRNA decay.

Across the host genome, Pol II appears to be most strongly affected at the stage of recruitment. The reduction in promoter-proximal Pol II is propagated into the gene body, although the differences there were less pronounced, suggesting that once transcription of a gene initiates there are not subsequent blocks to elongation. This agrees with prior studies of individual host loci in MHV68 infected or muSOX expressing cells (22,24). The population of Pol II that is promoter-proximal and measured during a ChIP experiment has been shown to comprise a combination of actively engaged Pol II as well as Pol II transiently binding to the DNA. Thus, a large reduction in promoter-proximal Pol II may not be expected to be borne out in the gene body (32). Additionally, the antibody we use in this study binds with higher affinity to hypophosphorylated Pol II(33), likely affecting our ability to pull down elongating and phosphorylated forms of Pol II. We also did not observe a defect in transcription termination, or evidence of polymerases on the DNA after TTSs, suggesting that the gammaherpesvirus MHV68 does not induce the termination defect and run-on transcription of host genes recently described in HSV-1 (34).

Two pieces of evidence indicate that reduction of Pol II at host genes is unlikely due to sequestration of or competition for the transcription machinery by the viral DNA. First, infection with the mRNA decay mutant virus R443I, which replicates its viral genome to WT levels (6), does not induce the widespread reduction in Pol II recruitment to host promoters. Second, transfection of muSOX alone (or other broad acting viral endonucleases) in the absence of viral infection is sufficient to induce transcriptional repression at host promoters. Therefore, we instead favor the hypothesis that one or more cellular proteins released from degrading mRNA fragments triggers a signaling event that restricts transcription machinery occupancy of mammalian chromatin (22).

Reduced Pol II occupancy of the host genome during MHV68 infection parallels infection data from the alphaherpesvirus HSV-1. HSV-1 infection results in a genome-wide reduction in Pol II occupancy of host chromatin and reduced nascent RNA production of the majority of host transcripts (34-36). Furthermore, early data showed that while the endogenous mouse β-globin locus was transcriptionally repressed during HSV-1 infection, its expression was rescued upon integration of the β-globin gene into the HSV-1 genome (37,38). Experiments with gene-specific null viruses have implicated immediate early (IE) proteins in the HSV-1 transcriptional repression phenotype, including ICP27, which can accelerate mRNA turnover (36,39). Additionally, we have previously published, and confirm in this report, that transfection of HSV-1 vhs, another broad acting endoribonuclease, is sufficient to induce reduced Pol II occupancy of several host promoters (21). Thus, in addition to targeting host and viral mRNAs for cytoplasmic decay, both alpha and gammaherpesviruses restrict Pol II occupancy of the host genome, although additional work is needed to establish whether the underlying mechanism is similar.

During the lytic stage of herpesvirus infection, viral DNA is localized to replication compartments. These non-membrane bound structures are sites of viral genome replication, transcription, and packaging (40). They are enriched for many host and viral proteins involved in these processes, including chromatin modifying factors and Pol II (28-30). Replication compartments grow and coalesce over infection, excluding host chromatin, which becomes pushed to the nuclear periphery (25-27). We recently showed that muSOX and Xrn1-coordinated mRNA decay causes differential trafficking of many cellular RNA binding proteins between the cytoplasm and nucleus. Knockdown and overexpression studies suggested that at least one of these proteins, PABPC, is associated with transcriptional repression (22). Here, we detected PABPC in viral replication compartments of infected cell nuclei, indicating that selective exclusion of PABPC from these compartments is not the mechanism by which the viral genome escapes repression. Instead, we hypothesize that either the state of viral DNA or other protective features of the replication compartment confers escape from transcriptional repression.

Protection from mRNA decay-induced transcriptional repression was not conferred to non-MHV68 or viral promoters present on the latent viral episome, indicating that lytic phase viral DNA replication or replication compartment formation is key to this phenotype. The latent genomes of gammaherpesviruses including KSHV and Epstein Bar Virus (EBV) are heavily methylated and histone-rich, with the exception of promoters of genes expressed during latency, the latent origin of replication, and CTCF binding sites (41-43). Notably, during latency key lytic promoters like RTA contain both the activating H3K4me3 and the repressive H3K27me3 histone marks, representing a ‘poised’ state (44). Rapid changes shortly after lytic cycle induction result in hypomethylation and removal of repressive marks at early gene promoters (41,44). At later stages of infection, KSHV, EBV and HSV-1 genomes have been shown to be largely nucleosome free, consistent with histones not being packaged in virions (27,29,45). For gammaherpesviruses, this is the stage at which mRNA degradation is most pronounced. We therefore hypothesize that the ‘open’ state of viral DNA during lytic replication plays an important role in facilitating Pol II recruitment under conditions of accelerated cytoplasmic mRNA decay. Future work is geared towards exploring whether widespread mRNA decay alters cellular chromatin states and DNA accessibility and, if so, how this influences transcription factor recruitment.

In conclusion, decay-coupled transcriptional repression is a novel facet of viral manipulation of the cellular gene expression landscape. We hypothesize that the ability to escape from transcriptional repression is linked to the open conformation of viral DNA in replication compartments, and that this helps these viruses counteract decay-induced viral transcript loss. While widespread mRNA decay has been shown to facilitate immune evasion, gene expression control, and latency establishment, the robust viral transcription we document here presents a parsimonious explanation for a novel way in which the virus benefits from accelerated decay.

## Materials and Methods

### Plasmids and Primers

pCDEF3 muSOX and VHS and SOX have been previously published (24). pCDEF3-GFP was used as a control.

Primers used for qPCR and for cloning are listed in S2 Table. M7, M4 and CMV-puro were made by cloning the respective promoter sequence upstream of the puromycin cassette in a pLKO-sgRNA vector. The promoter of M4 was defined as the 1131bp upstream of the M4 coding region not overlapping with other viral ORFs and the promoter sequence of M7 was previously published (46). pGL3-kLANA-puromycin was made by introducing the puromycin gene into the I-CeuI and XbaI sites of pGL3-kLANA via InFusion (Clonetech) cloning. pGL3-kLANA was made by amplifying KSHV’s LANA promoter sequence (47) and using InFusion cloning to introduce it into the XhoI and HindIII sites of pGL3.

### Cells and transfections

MC57G mouse fibroblast cells (ATCC), 293T cells (ATCC) and NIH 3T3 cells (ATCC) were maintained in DMEM (Invitrogen) supplemented with 10% fetal bovine serum (FBS). iSLK BAC16 cells (48) were maintained in DMEM supplemented with 10% FBS and 1 mg/mL hygromycin B. To reactivate iSLK cells, they were treated with 1 μg/mL doxycycline and 1 mM sodium butyrate for 48 hours.

DNA transfections were carried out in 293T cells at 70-90% confluency in 10 cm plates with 10 μg host-shutoff factor plasmid or vector control using PolyJet (SignaGen) 24 hours before harvesting and again at 18 hours before harvesting.

Lentiviral transduction was carried out by spinfecting 1 × 10^6^ 293T or MC57G cells with lentivirus made from 2^nd^ generation plasmids at a MOI <1 for 2 hours with 4 μg/mL polybrene (Fisher). Twenty-four hours later puromycin (1 μg/ml for 293T or 3 μg/ml for MC57G) was added to select for transduced cells. 293T cells stably expressing the kLANA-puromycin cassette were made by transfecting the cells with 1 μg pGL3-kLANA-puromycin using PolyJet, then selecting with puromycin for random integration events.

For the CMV-luciferase promoter assay, populations of 293T cells expressing muSOX were selected 24 hours post-transfection using the Miltenyi Biotec MACS cell separation system according to the manufacturer’s instructions as previously described (22).

293T cells were infected with KSHV by reactivating iSLK-Bac16 cells for 48 hours and then spinfecting cell-free virus onto a monolayer of 293T cells, as described for lentiviral production but in the absence of polybrene.

3 million MC57G cells were nucleofected with a 100uL Neon tip (1400 Voltage/20 Width/2 Pulse) to knockdown Xrn1 using a pool of 4 siRNAs (Dharmacon M-046621-01) or a non-targeting control (Dharmacon D-001206-14) using 10 μL of 20 μM siRNA.

### 4SU labeling

Adapted from (49).Cells were incubated in suspension with 500 μM 4SU (Sigma T4509) in DMEM supplemented with 10% FBS for 10 min then washed with PBS. RNA was extracted with TRIzol followed by isopropanol purification. RNA (300 μg) was incubated in biotinylation buffer (50 mM HEPES [pH 7.5], 5 mM EDTA) and 5 μg MTSEA-biotin (Biotium #90066), rotating at room temperature in the dark for 30 min. A no biotin control was made from an equal amount of total RNA and incubated as above but without the addition of MTSEA-biotin. RNA was then phenol:chloroform extracted and precipitated with isopropanol. The pellet was resuspended in DEPC-treated water and mixed with 50 μL Dynabeads MyOne streptavidin C1 (Invitrogen) that had been pre-washed twice with wash buffer (100 mM Tris [pH 7.5], 10 mM EDTA, 1 M NaCl, 0.1% Tween 20). Samples were rotated in the dark for 1 hour at room temperature, then washed twice with 65°C wash buffer and twice with room temperature wash buffer. Samples were eluted twice with 100 μL 5% Beta-mercaptoethanol (BME) in DEPC H_2_0 for 5 min then RNA was precipitated with isopropanol and quantified by RT-qPCR.

### Viral mutagenesis, propagation and infections

The WT and R443I MHV68 bacterial artificial chromosome (BAC) were previously described (6,50). MHV68 was produced by first making p0 virus by transfecting NIH 3T3 cells in 6-well plates with 2.5 μg BAC DNA using TransIT-X2 (Mirus Bio) for 24h. Five to seven days later, the cells were split into a 10 cm dish, then harvested and frozen. After 5-7 days the majority of cells showed cytopathic effect (CPE). 30 μL of p0 was then added per confluent 10 cm dish of NIH 3T12 cells, split 2 days later into four 10 cm dishes and harvested 4-6 days later when all cells were infected. Cells were collected by centrifugation (5 min, 1500 rpm) and the pellet was dounced 10 times. The cell pellet and supernatant were then ultracentrifuged for 2 h at 30,000 rpm. The resulting pellet was resuspended and titered by plaque assay. Cells were infected with MHV68 at an MOI of 5 for 24 hours.

### Western blotting

Cells were lysed in RIPA buffer (50 mM Tris-HCl pH7.6, 150 mM NaCl, 3 mM MgCl_2_, 10% glycerol, 0.5% NP-40, cOmplete EDTA-free Protease Inhibitors [Roche]) and then clarified by centrifugation at 21,000 × g for 10 min at 4°C. Whole cell lysate was quantified by Bradford assay and resolved by SDS-PAGE. Antibodies for western blot are: Xrn1 (Bethyl A300-443A, 1:1000), and Gapdh (Abcam, ab8245, 1:2000).

### RT-qPCR

RNA was reverse transcribed using AMV RT (Promega) with random 9-mer primers. Total RNA was DNase treated with TURBO DNase (ThermoFisher) or DNase treated on column with Zymo Direct-zol RNA MiniPrep Plus DNase. cDNA was quantified using iTaq Universal SYBR Mastermix (BioRad) and transcript-specific primers. All qPCR results are normalized to 18S levels and WT or vector control set to 1.

### Immunofluoresence

NIH 3T3 cells were plated on coverslips (7.5 × 10^4^ cells/well of a 12-well), infected the following day for 25-27 h, and then fixed in 4% formaldehyde for 10 min. iSLK-Bac16 cells were plated on coverslips (5 × 10^4^ cells/well of a 12-well), reactivated 24h later with doxycycline and sodium butyrate for 48 hours and then fixed as above. Cells were permeabilized with ice-cold methanol at −20°C for at least 20 min and incubated with anti-Pol II antibody (Biolegend, 8WG16 at 1:200), anti-PABPC antibody (Abcam ab21060 at 1:200 for mouse cells and Santa Cruz sc32318 at 1:25 for human cells) or anti-ORF59 antibody (Advanced Biotechnologies 13-211-100 at 1:200) in 5% BSA overnight at 4 °C. Secondary antibodies were added (1:1000) for 1 h at 37 °C. Coverslips were mounted in DAPI-containing Vectashield (VectorLabs). We identified RCs by DAPI exclusion and Pol II recruitment or ORF59 staining (28,30,51). Images were collected on a Zeiss LSM 710 AxioObserver with a 40x oil objective. Cells with RCs were first identified, then PABPC co-localization was determined by counting cells with at least 2x nuclear pixel intensity relative to non-RC containing cells in the same image (ImageJ).

### Chromatin immunoprecipitation (ChIP)

ChIP was performed from 15 cm plates of cells. 3 × 10^6^ MC57G or 293T cells were plated 24 h before infection and then harvested 24 h after infection or transfection(s). Upon harvesting, cells were washed with PBS, crosslinked in 1% formaldehyde at room temperature for 10 min. Cells were quenched in 0.125 M glycine for 5 min and washed twice with cold PBS. Crosslinked cell pellets were mixed with 1 mL fractionation buffer (5mM PIPES pH 8.0, 85 mM KCl, 0.5% NP-40 with cOmplete EDTA-free Protease Inhibitors [Roche]) and incubated on ice for 10 min, during which time the lysate was dounce homogenized to release nuclei and spun at 4,000 rpm for 5 min at 4°C to pellet nuclei. Nuclei were then resuspended in 300 μl of nuclei lysis buffer (50mM Tris-HCl pH 8.0, 0.3% SDS, 10mM EDTA, with cOmplete EDTA-free Protease Inhibitors [Roche]) and rotated for 10 min at 4°C followed by sonication using a QSonica Ultrasonicator. Sonicated chromatin was then spun at 13,000 rpm for 10 min at 4°C and the pellet discarded. 40 μg of chromatin measured with the Qubit broad range DNA reagent (ThermoFisher) was brought to 500 μL with ChIP dilution buffer (16.7 mM Tris-HCl pH 8.0, 1.1% Triton X-100, 1.2 mM EDTA, 167 mM NaCl and cOmplete EDTA-free Protease Inhibitors [Roche]) and incubated with 10 μg mouse monoclonal anti-RNAPII (BioLegend, 8WG16) or mouse IgG (Fisher Scientific) overnight, whereupon samples were rotated with 20 μl protein G dynabeads (Thermofisher) for 2 hours at 4°C. Beads were washed with low salt immune complex (20 mM Tris pH 8.0, 1% Triton-x-100, 2 mM EDTA, 150 mM NaCl, 0.1% SDS), high salt immune complex (20 mM Tris pH 8.0, 1% Triton-x-100, 2 mM EDTA, 500 mM NaCl, 0.1% SDS), lithium chloride immune complex (10 mM Tris pH 8.0, 0.25 M LiCl, 1% NP-40, 1% Deoxycholic acid, 1 mM EDTA), and Tris-EDTA for 5 min each at 4°C with rotation. DNA was eluted from the beads using 100 μl of elution buffer (150 mM NaCl, 50 μg/ml Proteinase K) and incubated at 55°C for 2 hours, then 65°C for 12 hours. DNA was purified using a Zymo Oligo Clean & Concentrator kit, and quantified by qPCR using primers to the promoter regions of indicated genes for 50 cycles. qPCR was simultaneously performed on input chromatin and normalized to input. Raw percent input values are normalized by setting mock infected or a transfection control to 1 and are plotted as relative % input. IgG pull downs were performed with the MHV68 infected sample under scramble conditions, or the muSOX or SOX transfected sample for transfection experiments.

### ChIP Sequencing and data analysis

Libraries were prepared for sequencing (Kapa Hyper Prep) using equal amounts of chromatin within each experiment, amplified for 10-12 cycles based on Kapa’s recommendations and sequenced on a HiSeq4000 with 100bp single end reads or 150bp reads (Xrn1kd second replicate only). Sequencing quality was assessed with FastQC and trimmed with Sickle. A custom index was made of the mm10 and MHV68 genomes with Bowtie2 build. Reads were mapped to that index using Bowtie2 (2.3.0).

To visualize reads in the Integrative Genomics Viewer (IGV), tdf files were made from bam files and indexes made from those bam files. To make tdf files for the viral genome, a chrom.size file was created for MHV68 and then the count command was run. Analysis of mapped reads was performed with HOMER (Hypergeometric Optimization of Motif EnRichment) (52). Tag directories were created from the bam files generated from Bowtie with the makeTagDirectory command. This was done separately for the mouse and MHV68 genomes. For mouse genes, the analyzeRepeats command was used to look at tag density in the body of genes and the –c pausing option was used to compare pausing ratios. annotatePeaks was run in both the –hist and –ghist mode to assess composite Pol II occupancy across the mouse genome as well as individual gene occupancy.

Viral analyses were done by first creating a custom genome annotation in HOMER using the loadGenome command. The TSS file was created from data generously shared by Scott Tibbetts (University of Florida) including portions of the viral BAC provided by Laurie Krug (Stony Brook University). annotatePeaks was run in the –hist mode to assess composite Pol II occupancy across the viral genome as well as individual gene occupancy.

## Supporting information

Key Resources

## Acknowledgements

We thank all members of the Glaunsinger and Coscoy labs for their helpful suggestions and discussions. We thank David McSwiggen for his microscopy assistance, Jennifer Kugel for her guidance applying HOMER to the viral genome, Scott Tibbetts for sharing MHV68 TSS data and Laurie Krug for providing a KSHV BAC sequence. This work used the Vincent J. Coates Genomics Sequencing Laboratory at UC Berkeley.

## Supporting information

**S1 Fig.**
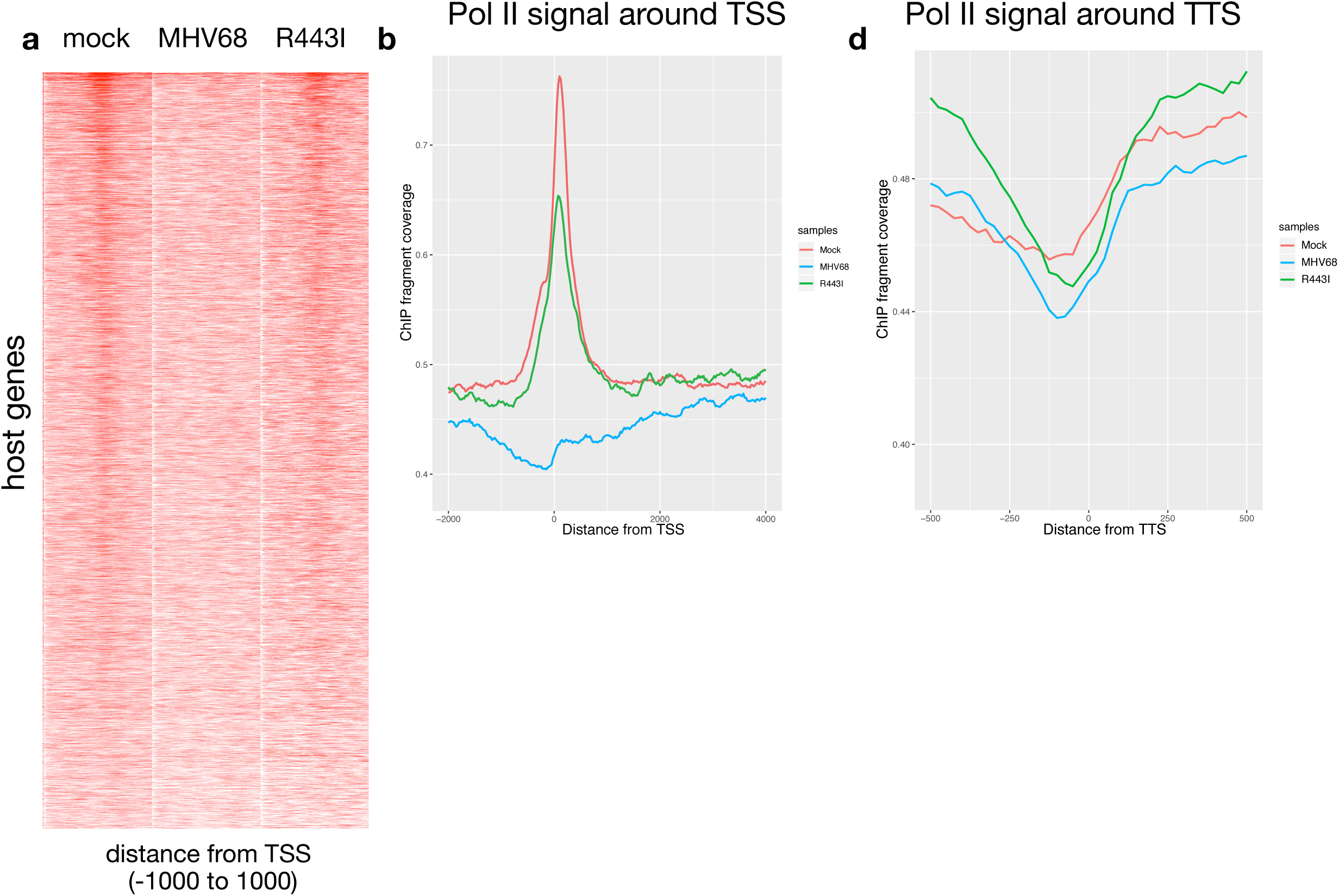
ChIP-seq replicate with WT and R443I MHV68. A) Pol II ChIP-seq signal profiles of host genes are shown in mock-infected, MHV68-infected and R443I-infected cells. Each row of the heat map displays Pol II occupancy of one gene from – 1000 to +1000 in 25 bp bins. Genes are ranked by the Pol II-transcription start site (TSS) proximal signal in mock infected cells. (B) Sequence tags were plotted as a histogram in 25 bp bins for −2000 to +4000 around the TSS. Mock, MHV68 (red) and MHV68 (blue) and R443I (green) traces are shown along with their input controls. (C) Pol II ChIP-seq coverage across the *Srsf2* gene for ChIP-seq replicate 1 and 2 with mock, WT MHV68 and R443I shown. Alignment files were converted to the tiled data file (tdf) format and visualized in the Integrative Genome Viewer. (D) Pol II transcription termination is not dependent on RNA decay. Sequence tags were plotted as a histogram in 25 bp bins for transcription termination sequence (TTS) proximal Pol II for −1000 to +1000 around the TTS with the same color scheme as (B).

**S2 Fig.**
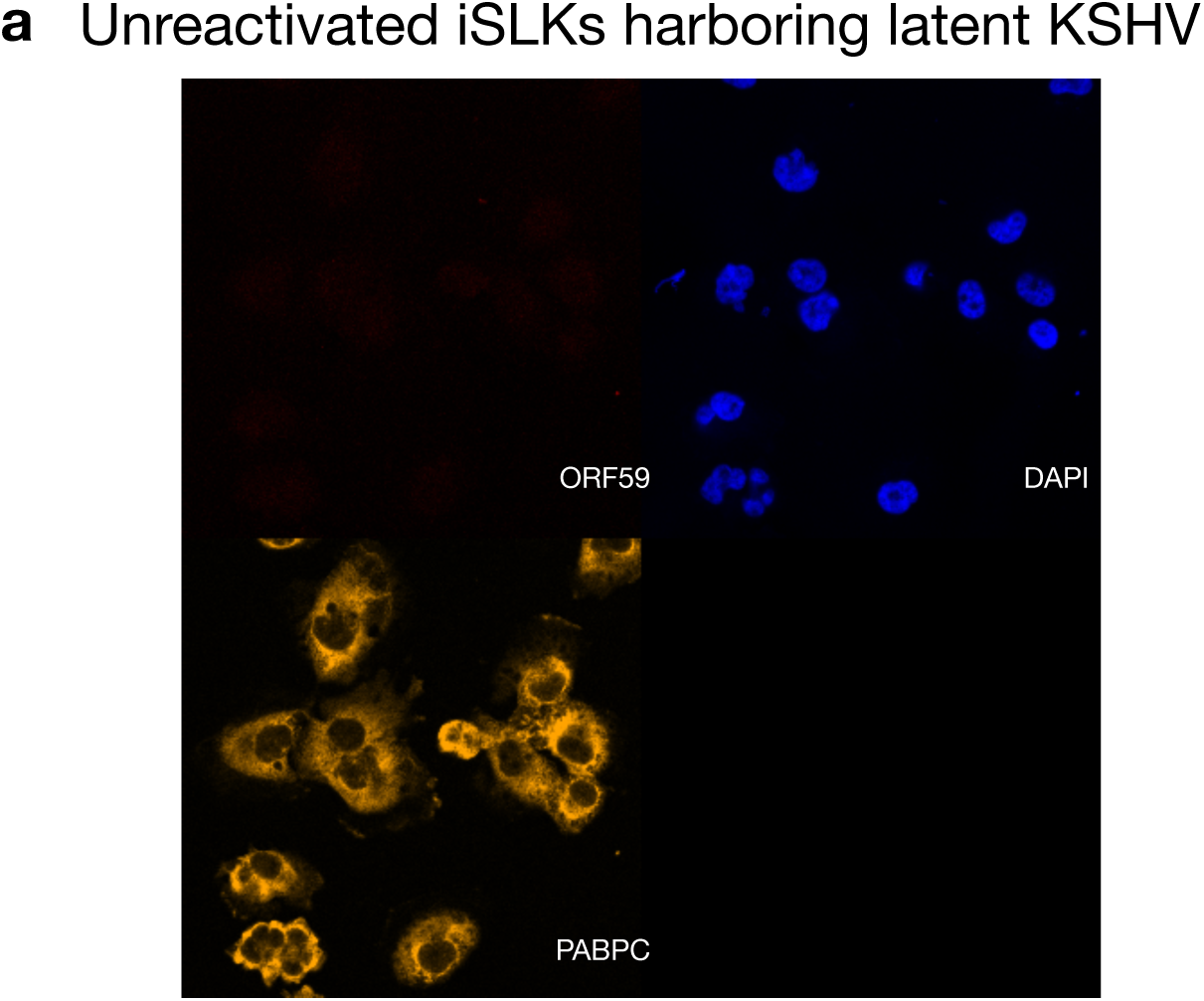
Unreactivated iSLK cells show primarily cytoplasmic PABPC signal. (A)IF was performed on unreactivated iSLK cells using antibodies recognizing PABPC and ORF59. Host DNA was stained with DAPI.

**S3? Fig.**
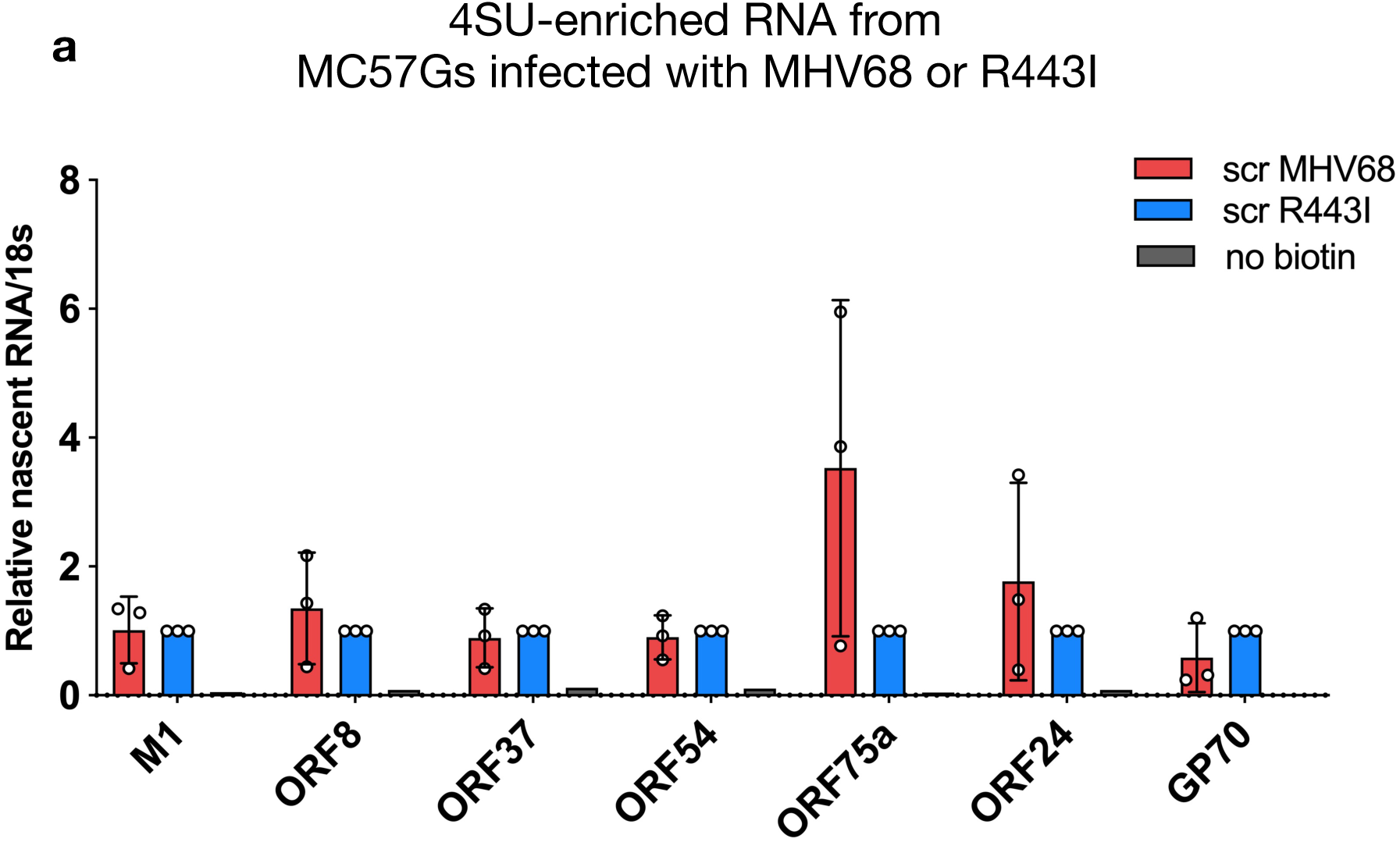
4SU-enriched RNA from MHV68 and R443I infected MC57G cells. (A) 4SU incorporated RNA levels of 7 viral genes in cells infected with MHV68 and R443I.

